# Prostaglandins and calprotectin are genetically and functionally linked to the Inflammatory Bowel Diseases

**DOI:** 10.1101/2022.04.06.487252

**Authors:** Mohamad Karaky, Gabrielle Boucher, Saraï Mola, Sylvain Foisy, Claudine Beauchamp, Marie-Eve Rivard, Melanie Burnette, Hugues Gosselin, iGenoMed Consortium, Alain Bitton, Guy Charron, Philippe Goyette, John D. Rioux

**Affiliations:** Montreal Heart Institute Research Center, Montreal, Quebec, Canada; McGill University Health Centre, Division of Gastroenterology, Montreal, Quebec, Canada; Université de Montréal, Faculty of Medicine, Montreal, Quebec, Canada

## Abstract

**Background:** Genome wide association studies (GWAS) have identified and validated more than 200 genomic loci associated with the inflammatory bowel disease (IBD), although for most the causal gene remains unknown. Given the importance of myeloid cells in IBD pathogenesis, the current study aimed to uncover the role of genes within IBD genetic loci that are endogenously expressed in this cell lineage.

**Methods:** The open reading frames (ORF) of 43 genes from IBD-associated loci were expressed via lentiviral transfer in the THP-1 model of human monocytes and the impact of each of these on the cell’s transcriptome was analyzed using a novel RNA sequencing-based approach. We used a combination of genetic and pharmacologic approaches to validate our findings in the THP-1 line with further validation in human induced pluripotent stem cell (hiPSC)-derived-monocytes.

**Results:** This functional genomics screen provided evidence that genes in four IBD GWAS loci (*PTGIR, ZBTB40, SLC39A11* and *NFKB1*) are involved in controlling *S100A8* and *S100A9* genes expression, which encode the two subunits of calprotectin (CP). We demonstrated that increasing PTGIR expression and/or stimulating *PTGIR* signaling resulted in increased CP expression in THP-1. This was further validated in hiPSC-derived monocytes. Conversely, knocking-down PTGIR endogenous expression and/or inhibiting *PTGIR* signaling led to decreased CP expression. These analyses were extended to the known IBD gene *PTGER4*, whereby its specific agonist also led to increased CP expression. Furthermore, we demonstrated that the *PTGIR* and *PTGER4* mediated control of CP expression was dependent on signaling via adenylate cyclase and *STAT3*. Finally, we demonstrated that LPS-mediated increases in CP expression could be potentiated by agonists of *PTGIR* and *PTGER4*, or diminished by their antagonists, with an opposite effect on TNF-alpha expression.

**Conclusion:** Our results support a causal role for the *PTGIR, ZBTB40, SLC39A11* and *NFKB1* genes in IBD, with all four genes regulating the expression of CP in myeloid cells, as well as an important role for the prostacyclin/prostaglandin biogenesis and signaling pathways in IBD susceptibility and pathogenesis.

**Author Summary:** Crohn’s Disease and Ulcerative colitis are the two main types of inflammatory bowel diseases (IBD). These are debilitating chronic inflammatory diseases of the digestive tract. IBD pathogenesis is complex and involves multiple different cell types within the intestinal mucosa. While over 200 regions of the genome have been associated with susceptibility to IBD, for most the causal gene remains to be identified. In the current study we have focused on genes from IBD loci that are endogenously expressed in monocytes or macrophages, given the importance of these cells in IBD pathogenesis. Specifically, we modulated the expression of 43 genes from within validated IBD loci, in a human monocyte/macrophage cell line, and determined the impact of this increased expression on the rest of the transcriptome. We found evidence that four of these genes (*PTGIR, ZBTB40, SLC39A11* and *NFKB1*) control the expression of calprotectin, which is a proinflammatory molecule that is used as a marker of intestinal mucosal inflammation. We then elucidated how prostaglandin signaling via prostaglandin receptors PTGIR and PTGER4, another IBD gene, regulates calprotectin expression. This work provides evidence that all five genes are causal and provide the first link between calprotectin and disease susceptibility.

## Introduction

Inflammatory bowel diseases (IBD) are chronic inflammatory diseases of the digestive system. Crohn’s disease and ulcerative colitis are the two most common subtypes of IBD (1). IBD is a global disease with increasing prevalence, estimated to reach four million patients in North America by 2030 (2). While the etiology of IBD is still not fully known, there is growing evidence suggesting that there is a combination of genetic and environmental risk factors impacting on disease susceptibility, with the latter including gut microbial components that chronically stimulate the gastrointestinal immune system (3).To this point, it is clear that the innate and adaptive immune systems are both implicated in IBD pathology. For example, in the innate immune system, neutrophil infiltration and activation have been correlated with UC severity (4). CD14^+^ CD16^+^ monocyte infiltrates in the inflamed mucosa have been identified as a major proinflammatory immune cell population in CD (5). Furthermore, in IBD, macrophages massively infiltrate the intestinal mucosa (6) and are considered important effectors of pathology, producing inflammatory mediators such as TNF-alpha, IL1, IL6, and nitric oxide (7). This is further supported by recent single cell RNA sequencing of intestinal tissues demonstrating an important enrichment of monocytes, inflammatory M1 macrophages, activated DCs and plasmacytoid DCs in inflamed tissues from various gut locations in IBD patients and has been correlated with disease severity, while in contrast, the anti-inflammatory M2 subset was diminished with severity (8).

In terms of genetic risk, > 200 genomic loci have been associated with IBD, CD or UC (9-11). Interestingly, a transcriptome-based analysis of the genes located within these IBD loci found an enrichment of expression within various immune cells (12). In addition, regulatory variants in IBD loci were active in different immune cells such as CD4+ T, CD8+ T, CD19+ B cells and CD14+ monocytes (13). While genetic and subsequent functional studies have identified a handful of causal genes from these loci that are believed to act primarily within cells of the monocyte/macrophage lineage (e.g *IRGM, LRRK2, CARD9*), no large-scale functional screen of IBD genes had been performed in this cellular context. Given this, additional approaches are necessary to resolve ∼95% of IBD loci. It is believed that most association signals at Genome Wide Association Studies (GWAS) loci could be explained by common regulatory variants that control the expression of one or more genes in disease relevant cell types (13, 14).

Given these observations, we propose that expression-based screens of genes within IBD loci can provide valuable information regarding a gene’s function within a specific cellular context and functionally link different genes through their shared impacts on the cell’s transcriptome. In the current study we have focused on genes from IBD loci that are endogenously expressed in monocytes or macrophages, given the importance of these cells in IBD pathogenesis. Specifically, we modulated the expression of 43 genes from within validated IBD loci and determined the impact of this increased expression on the rest of the transcriptome. This functional genomics screen has provided evidence that genes in four IBD loci (*PTGIR, ZBTB40, SLC39A11* and *NFKB1*) are involved in controlling the expression levels of *S100A8* and *S100A9*, the two subunits of CP, an important IBD biomarker for monitoring disease severity(15, 16). Given this, we explored the effect of the prostaglandin (PG) receptors *PTGIR* and *PTGER4* associated with IBD (17), receptors of PG I2 and PG E2 respectively, on regulating the expression of CP in THP-1 cells and hiPSC-derived monocytes in order to elucidate the role of the PG pathway in IBD pathology.

## RESULTS

### Functional genomics screen of 43 IBD genes in the human THP-1 monocytic line

To better understand the functional impacts of IBD genes in disease susceptibility within the monocyte/macrophage cell compartment, we performed an expression-based screen of genes from known IBD loci. In order to define the genes to include in this transcriptomic screen, we began with the 167 IBD-associated loci that had been identified prior to the initiation of the current study (12) and prioritized 65 genes with endogenous expression profiles consistent with having a role in monocyte/macrophage functions. Of these, we successfully cloned and stably transduced 43 IBD-ORFs into the THP-1 monocytic cell model, each via three independent infections (**Table 1 in S1 Table**). The RNA from these stable cultures was analysed by bulk RNA sequencing. Following normalisation of this transcriptomic data, merging of samples from different experimental batches and merging of experimental replicates, the induction level (**Table 1 in S1 Table**) and impact on the cell’s transcriptome of each IBD-ORF was assessed. To do so, we determined the variance in expression of all detectable genes in the transcriptome across all the samples included in these analyses, and then determined the set of genes that increased or decreased in response to the expression each ORF – these “HITS” were defined as genes where the fold effect computed from the combined replicates was larger than two and the expression was outside the expected range of variation (**Fig 1A & Table 2 in S1 Table**).

**Fig 1.**
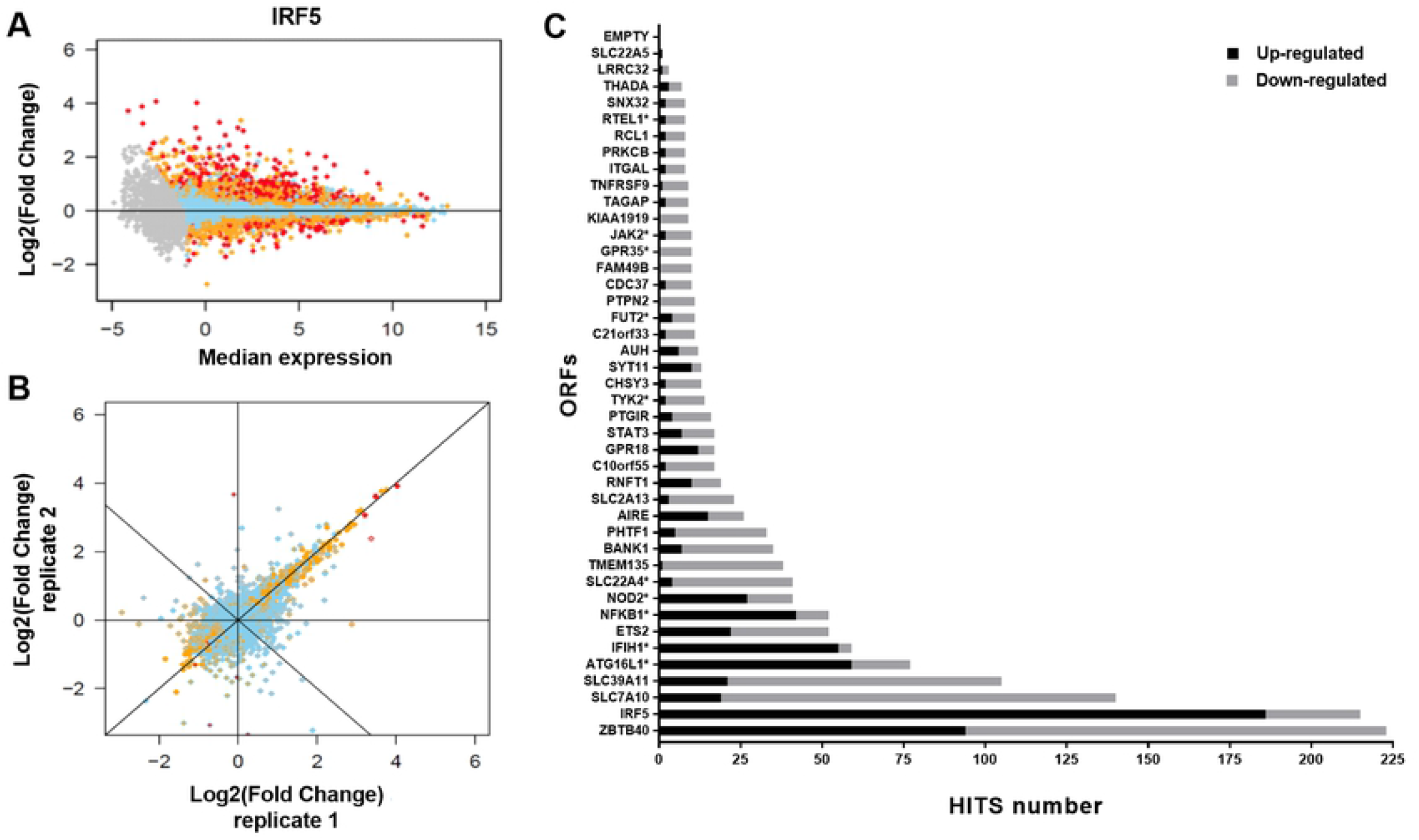
Impact of IBD gene candidate ORFs on the THP-1 transcriptome. **(A)** Selected example illustrating impact observed on the transcriptome of THP-1 cells following the expression of IRF5. Each dot represents a single detectable gene in the THP-1 transcriptome. The x-axis shows the log_2_-transformed median expression across all conditions tested (baseline). The y-axis represents the effect of transduction and expression of a given ORF, as the log_2_-transformed fold-induction compared to baseline. Skyblue dots represent genes with expression value within expected variation (|Z|≤2), orange dots represent genes suggestively outside the range (|Z|>2) and red dots represent genes outside expected range of variation (|Z|>4). Gray dots are genes with expression value below our detection threshold. **(B)** Correlation of effect of independent set of replicated expression of IRF5 on THP-1 transcriptome. The x-axis (inner color of dots) and y-axis (border color of dots) show the effect of two independent set of replicated ORFs on the transcriptome, as the log_2_-transformed fold-induction compared to baseline. Variation between sets of replicates includes effect of independent infection dates, RNA extraction, expression arrays and batches. **(C)** Impact of the transduction and expression of all 43 IBD gene candidate ORFs on the transcriptome of THP-1 cells. ORFs are ordered by their total number of HITS, with the number of up- and down-regulated HITS illustrated by black and gray, respectively **(Table 2 in S1 Table & S1 Appendix**). Starred ORFs are previously reported IBD candidate causal genes.

Importantly, there was strong correlation between the replicates of any given ORF, highlighting the robustness of the experimental approach used **(Fig 1B)**. We observed that the different IBD-ORFs showed a wide range of effects on the cell’s transcriptome (*from 1 to 223 HITS*), with IBD-ORFs encoding for known transcription factors (e.g. *ZBTB40, IRF5* and *IFIH1*) showing some of the greatest impacts on the transcriptome. On the other hand, ORFs encoding terminal enzymes in a metabolic pathway or proteins whose function likely requires an external stimulus, such as for the known causal genes *FUT2* (11 HITS) or *TYK2* (14 HITS), respectively, only had modest impacts on the transcriptome (**Fig 1C, Table 2 in S1 Table & S1 Appendix**).

### Expression of *IRF5* has a major impact on the THP-1 transcriptome

As a first step to interpreting the results from this ORF-based expression screen, we examined the results obtained for the ORF of interferon regulatory factor 5 (*IRF5*) as it is a gene that encodes a known transcription factor belonging to the interferon regulatory factor family of genes that are highly expressed in human monocytes, macrophages, dendritic cells and B cells (21). IRF5 plays a central role in inflammation by inducing the production of proinflammatory cytokines and promotes M1 macrophage polarization by directly inducing the transcription of M1 genes (22).

In this screen, the lentiviral transduction of the *IRF5* ORF increased the overall expression of this gene in THP-1 cells by ∼3-fold and resulted in the highest number of upregulated HITS (n=186) **(Fig 1 & Table 1 in S1 Table)**. This increased expression of *IRF5* resulted in a ∼10-fold or greater increase in the expression of proinflammatory mediators that belong to C-C motif chemokine ligand family (*CCL4, CCL4L1, CCL8*, and *CCL3*) and of the genes encoding the receptor for the proinflammatory cytokine IL18 (*IL18RAP* and *IL18R1*) **(Table 2 in S1 Table**). Notably, *CCL4* is also known to be upregulated and secreted by M1 macrophages. In addition, the increase in *IRF5* expression induced the expression of other M1 macrophages genes, such as the inflammatory cytokine *IL1B* and surface proteins CD83 and *CD40* implicated in antigen presentation and T cell activation, respectively **(Table 2 in S1 Table**) (22).

A global annotation analysis of the 186 genes that were upregulated following the increased expression of *IRF5* found significant enrichment of multiple terms, most of which are known to be implicated in the function of monocytic and other myeloid cells, including “defense response”, “innate immune response”, “response to cytokine” and “response to virus” (**Table 3 in S1 Table**). In addition, the analysis of the proximal promoters of these 186 upregulated genes, by two different tools (gProfiler and PRIMA), revealed significant enrichment for different IRF transcription factor binding sites (TFBS) **(Table 4 & 5 in S1 Table**), including *IRF5* itself and one of its HITS, *IRF7*. A significant enrichment for *TFBS* of the *STAT1*::*STAT2* heterodimer was also observed. *STAT1* is also a HIT upregulated by *IRF5*. Taken together, these results support that our experimental approach has the capacity to identify ORF-related functions relevant to myeloid biology.

### Multiple IBD genes are linked to the calprotectin pathway

As can be seen in **Fig 1C**, the *ZBTB40* ORF had a greater impact on the THP-1 transcriptome than *IRF5*, although its effect was nearly evenly split between positive (94 HITS with increased expression levels) and negative (129 decreased) effects. The function of *ZBTB40* has not been fully established, although consistent with the results from our screen, BTB-ZF proteins can act as transcriptional activators or repressors (23). A global annotation analysis of the 94 HITS that increased following the expression of *ZBTB40* in THP-1 indicated significant enrichment of multiple annotation terms most of them are known to play essential roles in myeloid cells such as: “myeloid leukocyte activation”, “neutrophil activation”, “Toll-like receptor binding” and “myeloid cell activation involved in immune response” **(Table 3 in S1 Table**). Consistent with this, the *S100A8* and *S100A9* genes that encode the two protein subunits that form CP were among the top 10 genes upregulated by *ZBTB40*. CP is a protein secreted by monocyte/macrophages and neutrophils, known to modulate the inflammatory response by stimulating leukocyte recruitment and inducing cytokine secretion (24). While there is widespread use of fecal CP as a biomarker of disease severity and mucosal healing in IBD, its role in susceptibility to IBD has not been previously reported(15). We therefore wanted to validate the potential link between the *ZBTB40* gene and *S100A8* and *S100A9* expression and thus performed an independent set of three transfections of THP-1 with the ORF of *ZBTB40* and observed a consistent increase in expression of *S100A8* and *S100A9* expression (**S2 Fig**).

Next, we examined our expression screen results for all other IBD-ORFs tested and found that, in addition to *ZBTB40*, the *NFKB1, SLC39A11* and *PTGIR* ORFs also led to an increase in expression of the CP *S100A8/A9* genes **(Table 2 in S1 Table & S1 Appendix**). Moreover, in the global annotation analyses of the HITS we found multiple annotation terms that were enriched and shared across these four IBD genes, such as: “calprotectin heterotetramer”, “iNOS-S100A8/A9 complex” and “neutrophil aggregation”, as well as “Toll-like receptor binding” shared between *PTGIR, SLC39A11* or *ZBTB40* **(Table 3 in S1 Table**). *NFKB1* is a transcription factor implicated in many biological processes such as inflammation, immunity, differentiation, cell growth, tumorigenesis and apoptosis. *NFKB1* controls the balance in the activation of pro-inflammatory and anti-inflammatory signaling pathways in the gut (25). *SLC39A11* encodes a zinc transporter that plays a crucial role in the Zn homeostasis, which is necessary for the innate immune system, especially for maintaining the function of macrophages(26). Finally, *PTGIR* encodes the receptor for prostacyclin (PGI2). While PGI2 has primarily been studied for the treatment of pulmonary hypertension (PAH), due to its effects on smooth muscle relaxation, more recent studies have revealed anti-inflammatory effects as well (27). Given the availability of agonists and antagonists for *PTGIR*, we focused our functional studies on this pathway.

As a first step, we performed an independent set of three transfections of the THP-1 line with the ORF of *PTGIR*. This resulted in an important induction in the RNA expression levels of both *S100A8* and *S100A9* **(Fig 2A)**, as originally observed in the transcriptomic screen. Conversely, knocking down the endogenous expression of *PTGIR* in THP-1 cells by approximately 80% led to a 50% reduction in the RNA expression levels of *S100A8* and *S100A9* (**Fig 2B**). Next, we evaluated the impact of *PTGIR* on CP protein expression. Specifically, we observed significant increases in CP levels in cell lysates (FC=6.7; *P*=1.17×10^−3^) and secreted into the culture supernatants (FC=2.9; *P*=1.11×10^−3^) in THP-1 following expression of the *PTGIR* ORF (**S3 Fig**). Together this confirms that *PTGIR* levels impact the levels of CP in THP-1 cells, and that modulation of the transcripts for *S100A8/A9* genes are accompanied with modulation of the protein concentration of S100A8/A9 dimer (CP).

**Fig 2.**
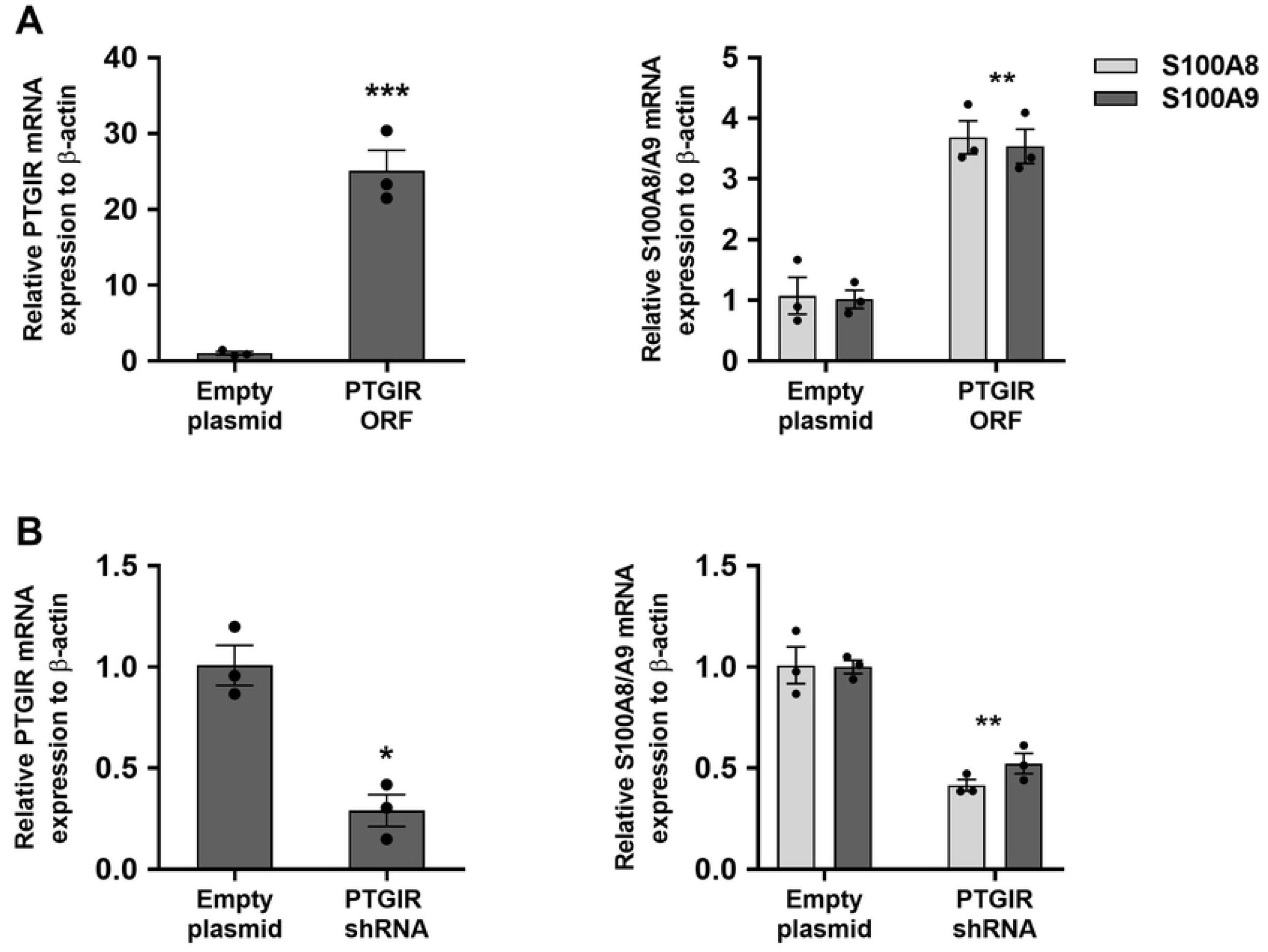
Impact of PTGIR expression and knockdown on S100A8/A9 genes expression in THP-1. **(A)** Relative mRNA expression levels of *PTGIR* and *S100A8/A9* genes in THP-1 cell lines following transduction with lentiviruses containing either an empty plasmid or a plasmid encoding for the *PTGIR* ORF. **(B)** Relative mRNA expression levels of *PTGIR* and *S100A8/A9* genes were evaluated in THP-1 cell lines following transduction with lentiviruses containing either an empty plasmid or a plasmid containing an shRNA targeting *PTGIR* (*PTGIR* shRNA). Each bar is the mean of 3 samples from 3 different infections ±SEM. \**P*<.05, \*\**P*<.01, \*\*\**P*<.001 (Student’s *t*-test paired).

### Genetic and pharmacologic modulation of *PTGIR* provides additional support for a link between *PTGIR* and CP pathways in monocytic cells

To provide additional support for this novel link between the *PTGIR* pathway and CP production, we studied the effects of well-characterized pharmacologic agents that act as a *PTGIR* agonist (Beraprost) or antagonist (Ro 1138452) on the expression of *S100A8*/*A9*. As can be seen in **Fig 3A**, endogenous levels of *PTGIR* are sufficient for THP-1 cells to respond to the *PTGIR* agonist Beraprost, as measured by the induction of *S100A8* (FC=6.25, *P*=9.98×10^−4^) and *S100A9* (FC=6.85, *P*=8.24×10^−5^) expression. This response is dramatically increased in THP-1 cells transduced with the *PTGIR* ORF (*S100A8* (FC=15.75, *P*=2.18×10^−5^) and *S100A9* (FC=20.46, *P*=2.49×10^−4^)) (**Fig 3B**) and is significantly decreased in cells where the expression of the endogenous *PTGIR* has been knocked down (*S100A8* (FC=2.47, *P*=.02) and S100A9 (FC=2.43, *P*=.04)) (**Fig 3C**). Furthermore, we found that the increase in *S100A8*/*S100A9* transcript following the stimulation of THP-1 parental cells with Beraprost was accompanied with an increase in secretion of the CP protein in the culture supernatant (FC=1.67, *P*=.01) (**S3 Fig**). Conversely, the use of the *PTGIR* antagonist Ro 1138452 led to significant decreases of *S100A8* (FC=0.46, *P*=5.78×10^−3^) and *S100A9* (FC=0.48, *P*=1.67×10^−3^) expression in wildtype THP-1 cells (**Fig 3D**) as well as in THP-1 cells expressing the *PTGIR* ORF (**Fig 3E**). However, no significant change has been observed following treatment with Ro 1138452 in cells where the expression of the endogenous *PTGIR* has been knocked down (**Fig 3F**).

**Fig 3.**
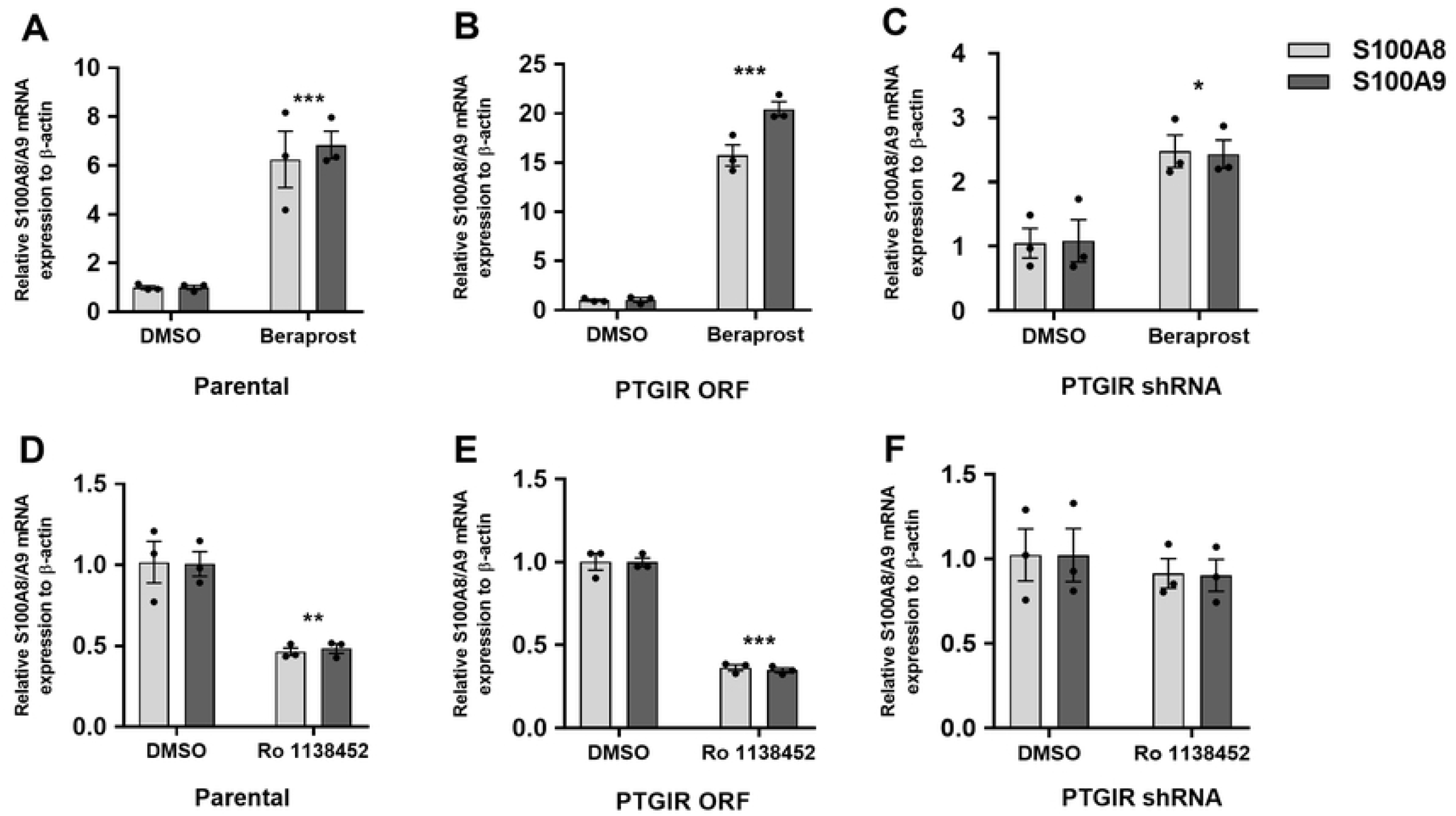
Impact of PTGIR agonist/antagonist on CP genes expression in THP-1: parental, PTGIR expressed or knocked-down. Relative mRNA expression levels of *S100A8/A9* genes were evaluated following the treatment of THP-1 parental cell line **(A)** with 10^−5^ M PTGIR agonist (Beraprost) or **(D)** with 10^−5^ M PTGIR antagonist (Ro 1138452) for 24 h. Relative in mRNA expression levels of S100A8/A9 genes were evaluated in THP-1 cell lines transduced with **(B, E)** PTGIR ORF or (**C, F**) PTGIR shRNA following a treatment with either DMSO and either 10^−5^ M PTGIR agonist (Beraprost) or 10^−5^ M antagonist (Ro 1138452) for 24 h. Each bar is the mean of 3 samples from 3 different infections ±SEM. \**P*<.05, \*\*\**P*<.001 (Student’s *t*-test paired). A dose and time-course response of *S100A8/A9* RNA expression to PTGIR agonist Beraprost were performed in parental THP-1 cells (**S7 Fig and S8, respectively**); A dose response of *S100A8/A9* RNA expression to PTGIR antagonist Ro 1138452 was performed in THP-1 cells transduced with the *PTGIR* ORF (**S9 Fig**).

### *PTGIR* signaling leads to transcriptional control of CP genes *S100A8*/*A9* via adenylyl cyclase dependent STAT3 signaling

It is known that *PTGIR* is a seven-transmembrane G-protein coupled receptor (GPCR) that is coupled to Gαs and that once this receptor is activated by its ligand, PGI2, the Gαs activates the adenylyl cyclase (AC), which converts the GTP into cAMP(28). In turn, the cAMP activates PKA, which then stimulates the activity of transcription factors such as CtBP1(29), SPI(30) and STAT3(31) by phosphorylation. As *S100A8/A9* appeared to be co-regulated in our dataset and in other studies (32), we evaluated their proximal promoters for the presence of shared TFBS. Specifically, using the publicly available ENCODE chip-seq data, we identified seven different TFs (CTCF, EP300, FOS, POLR2A, REST, SPI1 and STAT3) that had evidence of binding to both the *S100A8/A9* promoters (**S4 Fig**). Of these seven, only STAT3 is known to be activated via the cAMP-PKA pathway and thus constituted our best candidate (33). Given that transcription control mechanisms can vary from one cell type to another, and that the ENCODE ChIP-seq data for STAT3 was generated using the human epithelial cell MCF10A, we sought to validate this candidate pathway in THP-1 cells. As a first step, we studied the impact of the addition of 10µM Forskolin, a known activator of AC, on the induction of *S100A8/A9* expression in parental THP-1 cells. We found that this activation of AC led to a pronounced increase in expression of both *S100A8* (FC=15, *P*=1.41×10^−3^) and *S100A9* (FC=19, *P*=9.16×10^−4^), which was potentiated by the addition of the *PTGIR* agonist Beraprost (S100A8, FC=47, *P*=2.85×10^−4^; S100A9 FC=41, *P*=1.13×10^−5^) (**Fig 4A**). In contrast, we found no induction of *S100A8* or *S100A9* gene expression in cells treated by the AC inhibitor MDL12330A, in the presence of Beraprost (**Fig 4A**). Finally, we tested the role of STAT3 in this signaling pathway by treating the THP-1 parental cells with the STAT3 inhibitor Stattic.

**Fig 4.**
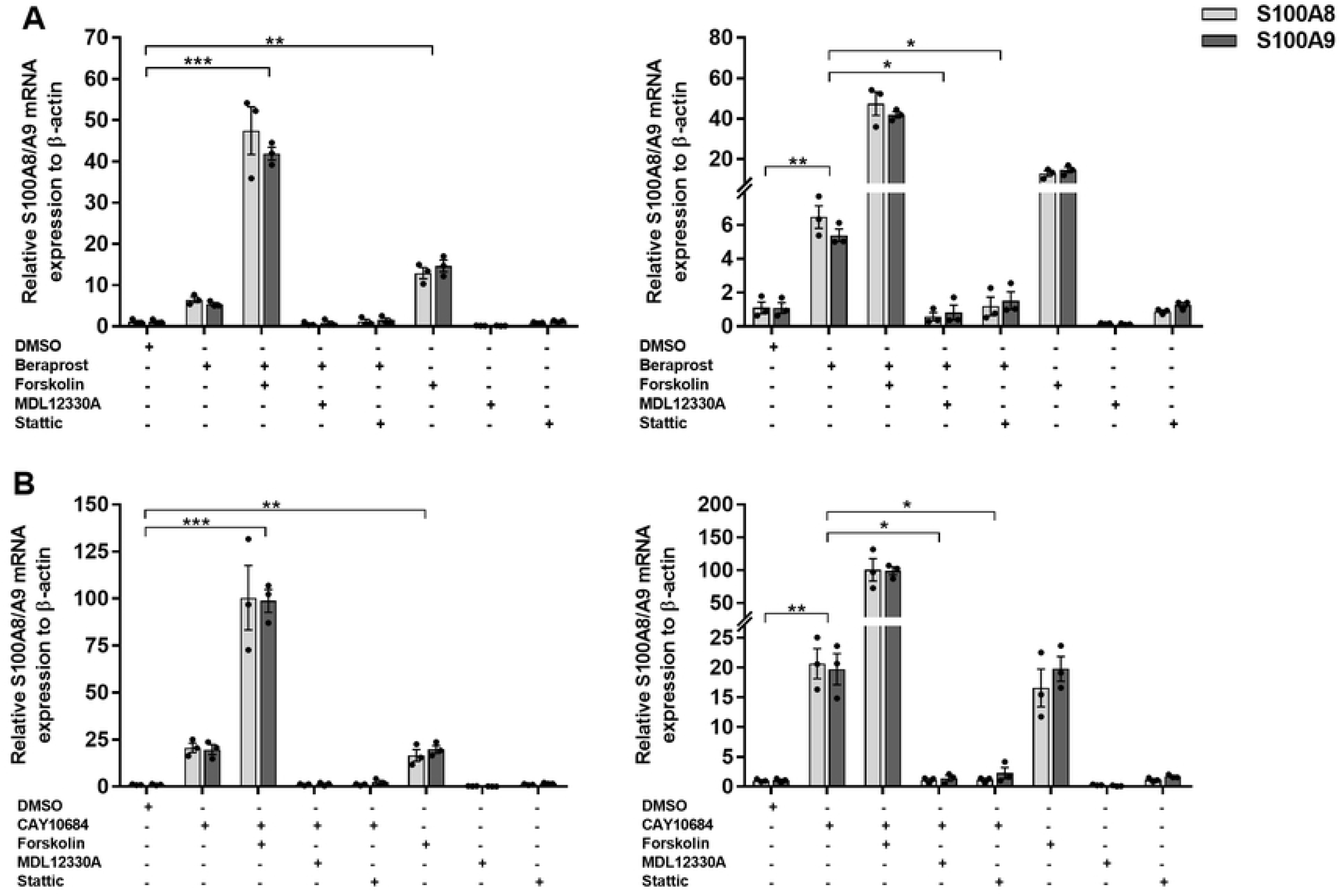
Impact of AC activator/inhibitor and STAT3 inhibitor on the *S100A8/A9* expression induction in parental THP-1. Relative mRNA expression levels of *S100A8/A9* genes were evaluated after treating 5×10^5^ THP-1 cells with either 10µM AC activator (Forskolin), 10µM AC inactivator (MDL12330A) or 2×10^−5^ M STAT3 inhibitor (Stattic) in the presence or absence of **(A)** 1×10^−5^ M PTGIR agonist (Beraprost) or **(B)** 1×10^−5^ M PTGER4 agonist (CAY10684) for 24h. Graphs on the right represents the same data with different y-axis scale. Each bar is the mean of 3 samples from 3 different experiments ±SEM. \**P*<.05, \*\**P*<.01, \*\*\**P*<.001 (Student’s *t*-test paired).

Specifically, treatment of the THP-1 cells with 2.10^−5^ M Stattic abrogated the response to the PTGIR agonist Beraprost (**Fig 4A**). These results validate that activation of the prostacyclin pathway leads to an increase in the expression of S100A8/A9 via an AC dependent STAT3 signaling pathway.

### The *PTGER4* pathway also regulates *S100A8/A9* expression

Based on our findings that activation of the PTGIR leads to increased expression of CP, as well as the findings that an increased expression of *ZBTB40* not only led to an induction of *S100A8/A9*, but also of three enzymes involved in PG pathway, *PLA2G1B, TBXAS1* and *AKR1C2* **(Table 2 in S1 Table)**, we were interested in investigating the involvement of other members of the PG pathway in the control of expression of *S100A8*/*A9*. In particular, the locus containing the *PTGER4* gene, which encodes the PG E2 receptor (subtype 4), was one of the first to be identified as a risk factor for CD, and then shown to be associated with both CD and UC phenotypes (12, 17) and is the predominant locus in genetic studies of IBD in African Americans (34, 35). Importantly, in the same study non-coding variants were found to be associated with increased expression level of *PTGER4* and were proposed as causal alleles, an observation that was later confirmed by others (17, 36). Thus, we decided to study the potential impact of PTGER4 signalling on *S100A8/A9* expression in the human myeloid model THP-1. As can be seen in **Fig 4B**, endogenous levels of *PTGER4* are sufficient for THP-1 cells to respond to the PTGER4-specific agonist CAY10684, as measured by the expression of *S100A8/A9*. Indeed, when these cells were exposed to CAY10684, we observed a very important induction in the expression of *S100A8* (FC=18.9, *P*=7.97×10^−5^) and *S100A9* (FC=20, *P*=2.16×10^−4^), which was equivalent to the induction of S100A8/A9 observed with the AC activator Forskolin. As was observed for PTGIR, the combination of the PTGER4 agonist and Forskolin led to a synergistic increase in *S100A8* (FC=100.3, *P*=3.33×10^−5^) and S100A9 (FC=98.7, *P*=1.97×10^−5^). Moreover, the induction of *S100A8/A9* by the PTGER4 agonist CAY10684 was abrogated by the addition of the AC inhibitor MDL12330 (**Fig 4B**). Finally, the induction of S100A8/A9 by CAY10684 was also abrogated by the addition of Stattic, the inhibitor of STAT3 (**Fig 4B**), suggesting a similar signaling pathway as was observed for PTGIR.

### The impact of PG receptors on *S100A8/A9* and on TNF-alpha in response to LPS

We also studied the effect of the PG receptor agonists on *S100A8/A9* in THP-1 stimulated by LPS. As can be seen in **Fig 5**, LPS induced the expression of *S100A8/A9* (*S100A8* FC=65.7 *P*=1.09×10^−3^, *S100A9* FC=81.2, *P*=1.38×10^−4^), and also induced the expression of PTGIR **(S5 Fig)**. In addition, PTGIR and PTGER4 agonists showed synergistic effect on S100A8/A9 expression in THP-1 when stimulated by LPS. Specifically, *S100A8/A9* was significantly induced when THP-1 cells were treated with the PTGIR agonist in the presence of LPS (S100A8: FC=2.39, *P*= .047; S100A9: FC=1.51, *P*=.014) or with the PTGER4 agonist in the presence of LPS (S100A8: FC=4.44, *P*=.032; S100A9: FC =2.39, *P*=.044) **(Fig 5)**. In contrast, when THP-1 cells were treated for 24h with Beraprost or CAY10684 (the agonists of PTGIR and PTGER4 respectively), there was a significant decrease in TNF-alpha induction in response to LPS stimulation (**Fig 6**). Specifically, there was almost a three-fold decrease in the amount of TNF-alpha induced in response to LPS in the presence of Beraprost (FC=0.38, *P*=.019), or CAY10684 (FC=0.30, *P*=2.9×10^−3^).

**Fig 5.**
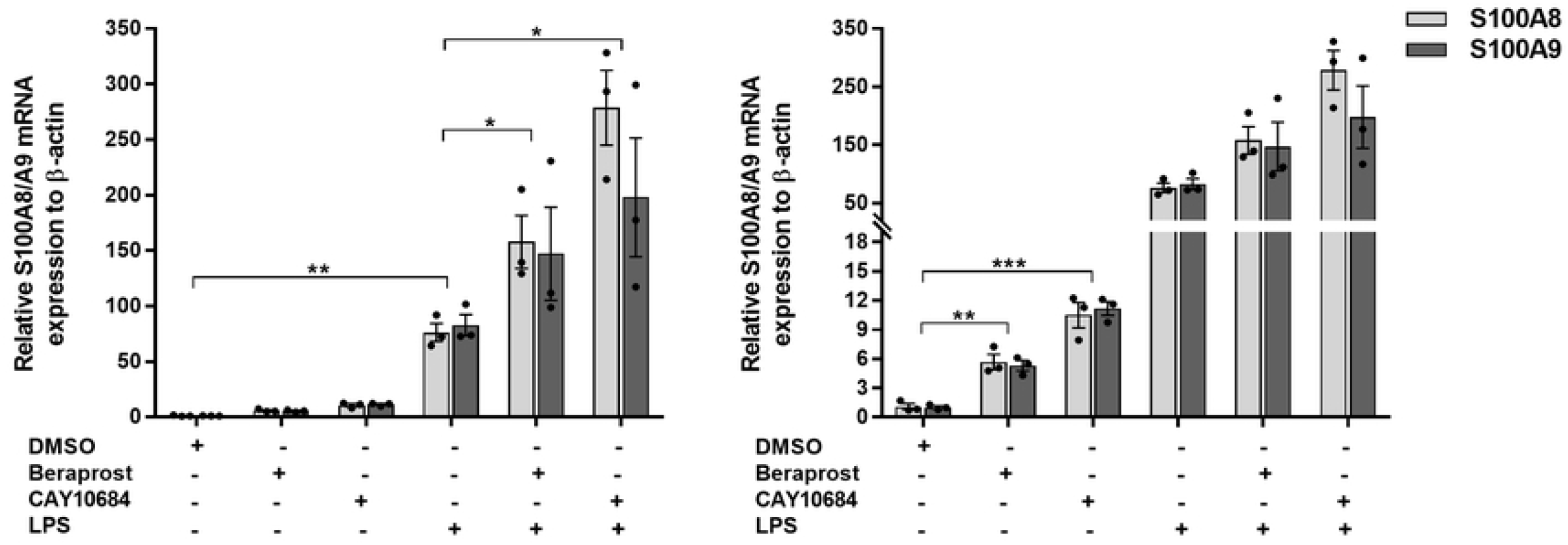
Impact of PG receptor agonists on the expression of *S100A8/A9* in response to LPS. Relative in mRNA expression levels of *S100A8/A9* genes were evaluated after incubating THP-1 for 24 hours with or without 0.2 ug/ml of LPS in the presence or absence of 1×10^−5^ M of Beraprost or CAY10684, the agonists of PTGIR and PTGER4 respectively. Graph on the right represents the same data with different y-axis scale. Each bar is the mean of 3 samples from 3 different experiments ±SEM. \**P*<.05, \*\**P* <.01, \*\*\**P*<.001 (Student’s *t*-test paired).

**Fig 6.**
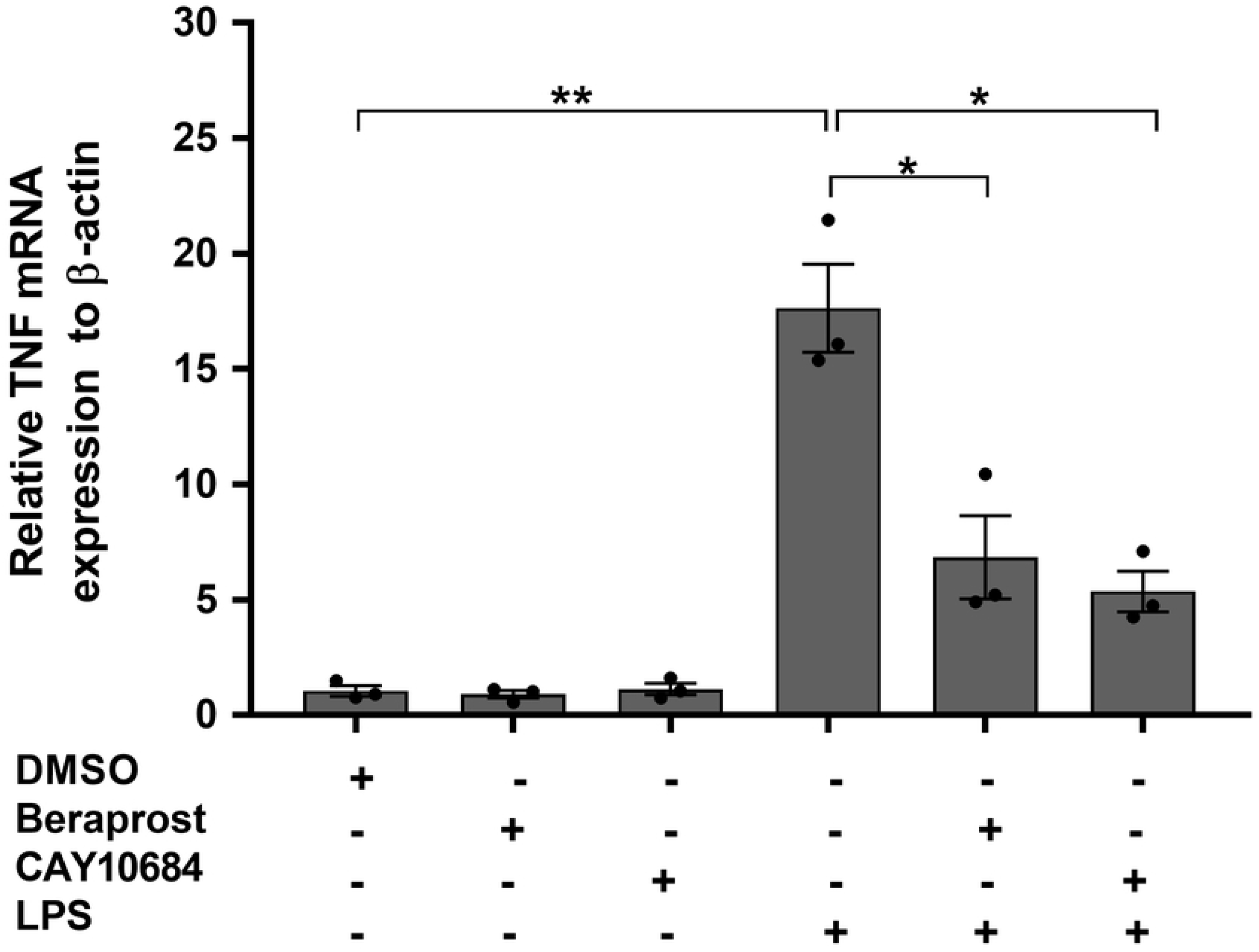
Impact of PG receptors on the inflammatory cytokine TNF-alpha in myeloid cells. Relative in mRNA expression levels of TNF gene were evaluated after incubating THP-1 for 24 hours with or without 0.2 ug/ml of LPS in the presence or absence of 1×10^−5^ M of Beraprost or CAY10684, the agonists of PTGIR and PTGER4 respectively. Each bar is the mean of 3 samples from 3 different experiments ±SEM. \**P*<.05, \*\**P*<.01 (Student’s *t*-test paired).

### Validating the role of prostaglandin pathways on the control of CP in human hiPSC-derived monocytes

Although THP-1 are widely used as a model of human monocytes, they derive from a patient with acute monocytic leukemia, thus to study the PG-CP connection in non-leukemic cells, we studied the impact of the PTGIR agonist on the expression of the CP genes in hiPSC-derived monocytes. Specifically, we found that the treatment of hiPSC monocytes with the PTGIR agonist Beraprost resulted in a significant induction of *S100A8* (FC=2.34, *P*=8.89×10^−3^) and *S100A9* (FC=1.74, *P*=.026) gene expression, providing additional support for this pathway **(Fig 7)**.

**Fig 7.**
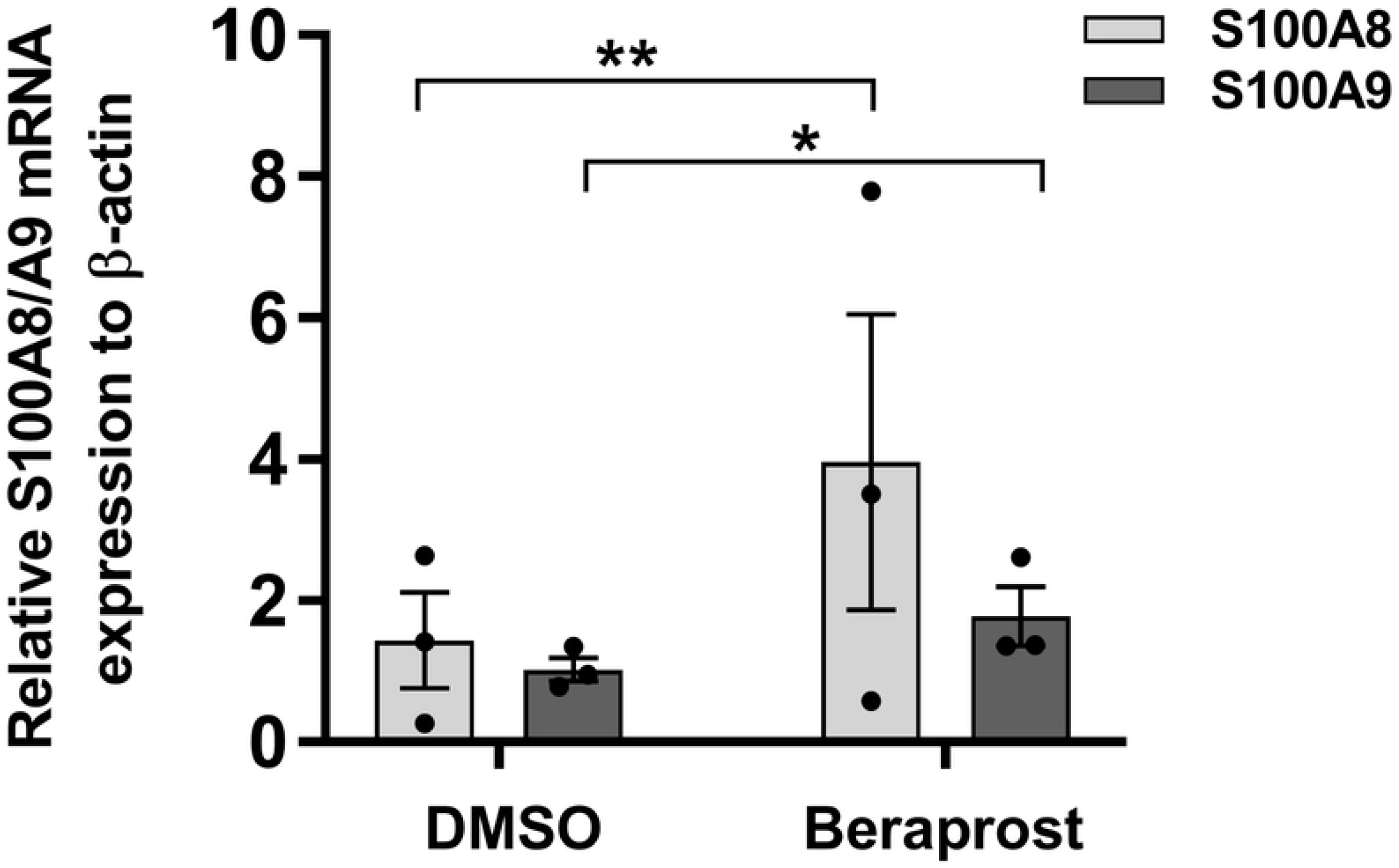
Impact of PTGIR agonist on the expression of CP genes in hiPSC-derived monocyte. **(A)** Relative in mRNA expression levels of *S100A8/A9* genes were evaluated following the treatment of hiPSC CD14+ with 10^−5^ M PTGIR agonist (Beraprost) for 24 hours. Each bar is the mean of 3 samples from 3 different experiments ±SEM. \**P*<.05, \*\**P*<.01 (Student’s *t*-test paired).

### Co-localization of IBD-associated SNPs and cis-eQTLs in CD14+ monocytes

Given that our ORF-based screen demonstrated that modulation of the expression levels of genes within IBD loci impacted the biology of THP-1 cells, we next examined whether the IBD-associated SNPs in the *ZBTB40, NFKB1, SLC39A11, PTGIR, PTGER4* and other GWAS loci could potentially impact on the expression of these genes in monocytes and/or macrophages. With this objective, we examined whether GWAS index SNPs, or any SNPs that were highly correlated with these, were previously identified in a genome-wide eQTL study of naive and stimulated monocytes (20). Specifically, we took the GWAS index SNPs for the 43 genes tested in the current study, as well as for *PTGER4*, and identified 1,695 SNPs that were highly correlated (r2 > 0.7) with these (**Table 7 in S1 Table)**. Next, we determined whether any of these were known eQTLs in human monocytes or macrophages (20). We found that 101 of these SNPs overlapped with the previously identified eQTLs in CD14+ cells, affecting the expression of 69 genes **(Table 8 in S1 Table**). In doing so, we found eQTLS for multiple causal IBD genes: *ATG16L1, TYK2, IFIH1, PTGER4*, and *PTGIR*. While this does support that many of the IBD-associated GWAS SNPs modulate the expression of these genes, the direction of effect for these is not always consistent with orthogonal data for these genes **(Table 8 in S1 Table**). For example, the major allele of the GWAS SNP associating *PTGER4* with IBD, is associated with increased disease risk and higher *PTGER4* expression in CD14+ monocytes activated by LPS or IFN for 24h. Increased susceptibility to colitis, however, is observed in wild-type mice administered an EP4-selective antagonist, while EP4-selective agonists are protective (37). For PTGIR, the minor allele of the GWAS SNP is associated with increased disease risk and lower expression of this receptor in CD14+ monocytes or macrophages. As presented above, selective agonists for both *PTGER4* and *PTGIR* increase the expression of CP.

## Discussion

Large-scale comprehensive collaborative studies have mapped out an important portion of the complex genetic architecture of susceptibility to CD and UC, with over 200 genomic loci being associated with CD, UC or both. These genetic findings have helped to identify key biological pathways involved in disease susceptibility ranging from autophagy to cytokine signaling in immune cells as well as the contributions of epithelial cells to anti-microbial control and intestinal barrier functions (38-42). Moreover, the relevance of these pathways is underpinned by the fact that many are targets of current therapies such as anti-integrins, anti-IL12/IL23 agents and JAK inhibitors. Given that the causal gene has been identified in only a fraction of the known IBD loci, as well as the importance of monocyte/macrophage lineage in IBD pathophysiology, we performed a functional screen of 43 genes from IBD loci in a human cell line often used as a model system for the study of human monocyte/macrophage functions. The scientific premise for the screen was that by modulating the expression of genes from IBD loci that have endogenous expression within this cell lineage, clues to their biological functions could be, at least in part, derived from their impact on the cells’ transcriptome and thus provide potential functional links to the pathogenesis of IBD.

Our analyses of IRF5, one of the 43 genes studied, illustrate this approach. Specifically, the transduction of the ORF for IRF5 in THP-1 cells led to an important increase in expression of genes associated with the polarization towards proinflammatory macrophages, consistent with its known functions(22). Previous fine mapping of the IRF5 locus defined a region that contained three potential causal genes (*KCP, IRF5* and *TNPO3*) (14), and we propose that our findings support *IRF5* as the causal gene within this locus. Moreover, *IRF5* has been associated with susceptibility to different inflammatory and autoimmune diseases including rheumatoid arthritis, systemic lupus erythematosus, multiple sclerosis, and Sjörgen’s syndrome (43). This is further supported by animal models where a reduction in *IRF5* expression attenuated colitis in mice, but also led to impaired clearance of intestinal pathogens, supporting a model where homeostasis requires a fine balance in the immune system’s response to gut flora (44).

The results described herein also revealed a connection between four genes (*ZBTB40, SLC39A11, NFKB1*, and *PTGIR*), located in four different IBD risk loci, and the expression of the *S100A8/A9* CP genes. This suggests that these are the causal genes within their respective GWAS loci. CP is a proinflammatory protein and its concentration in fecal and serum samples was shown to reflect IBD severity, although its utility as a biomarker still requires additional technical and clinical validation(45). Given this, we chose to further validate this link to IBD susceptibility genes. Specifically, we have demonstrated that the upregulation of *PTGIR* expression in THP-1 induces the expression of the *S100A8/A9* encoding for the two CP subunits and stimulates the secretion of CP protein. In addition, we showed that activating parental THP-1 cells with a PTGIR agonist increased *S100A8/A9* expression as well as the intracellular CP protein level, while a PTGIR antagonist resulted in reduced *S100A8/A9* RNA expression levels. Thus, PTGIR signaling has an important impact on CP RNA and protein expression, as well as on its secretion into the extracellular milieu. Moreover, we have validated the pathway by which the PTGIR induces the CP genes in our model. We have shown that the activation of the PGI2 receptor by its agonist induces the expression of *S100A8/A9* genes by activating AC, which is known to trigger cAMP-PKA pathway (46). PKA is known to activate STAT3 via phosphorylation (31). Activated STAT3 can then bind to the regulatory elements of *S100A8/A9* genes (47), inducing their expression. Although not part of the screen described herein, we extended our functional analyses to the PTGER4 receptor as it is part of the prostaglandin signaling pathway, albeit having a distinct ligand to PTGIR, and because it is a known causal IBD gene. Indeed, our results also provided support for the PGE2-PTGER4 pathway in the control of CP synthesis in monocyte/macrophages, which is consistent with studies in prostate epithelial cells (48). We further demonstrated that PGE2-PTGER4 signaling is also mediated by cAMP to PKA to STAT3 signaling, elucidating a common signaling pathway for two IBD genes and CP production (**Fig 8**). Although the THP-1 cell line is a widely used model for studies of macrophage differentiation, activation, and innate immune functions, we sought to validate our findings in hiPSC-derived monocytes as they are a renewable source of non-cancerous cells with normal ploidy. Again, we observed that stimulation of these with a PTGIR agonist resulted in an increased production of CP.

**Fig 8.**
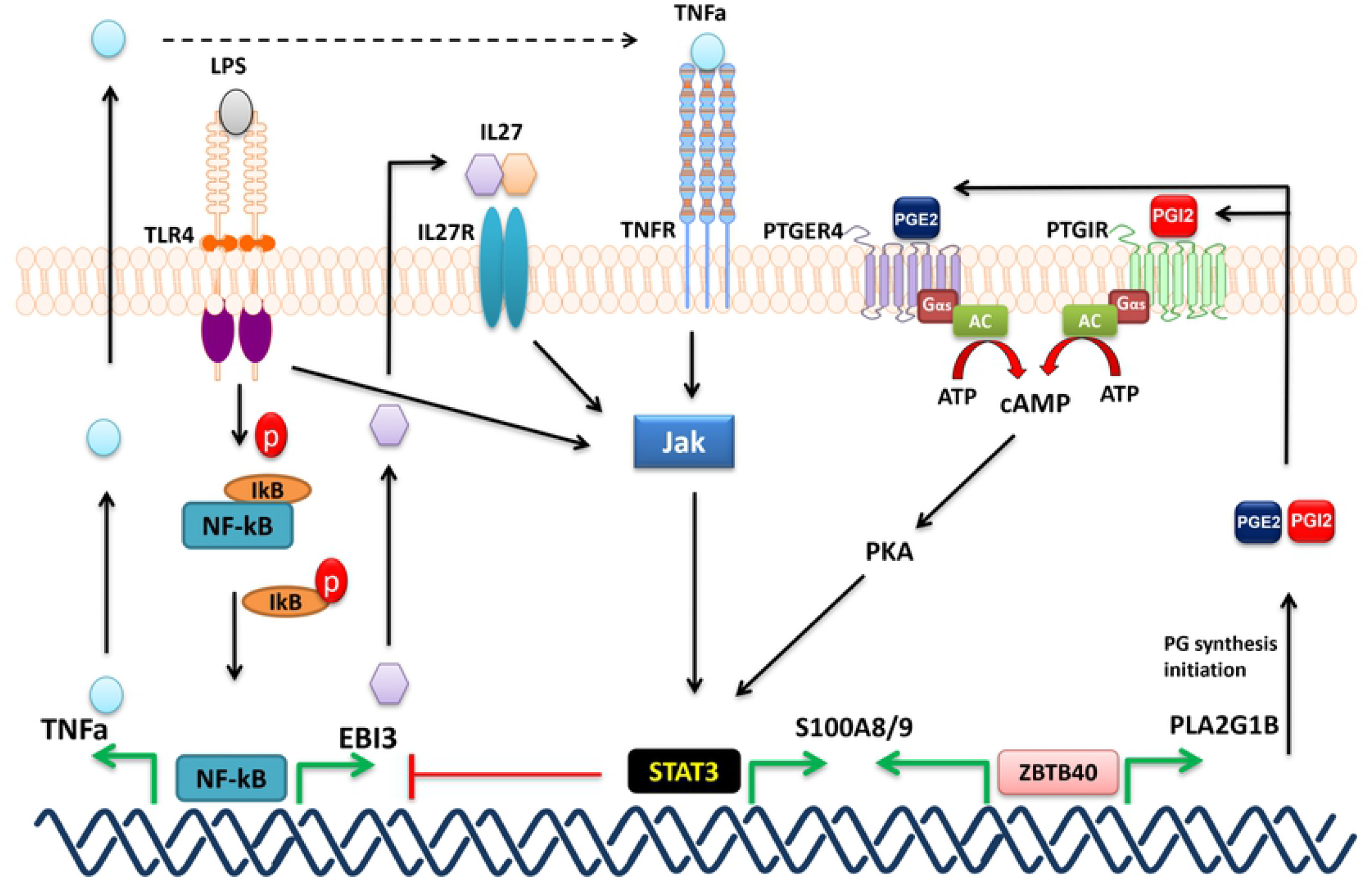
Proposed model of S100A8/9 induction. PTGIR and PTGER4 are activated by their ligands (PGI2 and PGE2 respectively) turning on the AC which converts ATP into cAMP; cAMP activates PKA which in turn triggers STAT3 by phosphorylation. STAT3 binds to the promoters (**S4 Fig**) of S100A8 and S100A9 inducing their expression. TLR4 and TNFR when activated by their respective ligands LPS and TNFa, activate the JAK2-STAT3 pathway which induces the expression of S100A8/9 and inhibits NF-kB. We have shown the induction of S100A8/9 genes in THP-1 activated by LPS **(Fig 5)** and TNF-alpha **(S6 Fig)**. ZBTB40 could potentially bind directly to the S100A8/9 promoters inducing their expression and/or to the PLA2G1B promoter (***by Encode***) activating its expression. PLA2G1B encodes an enzyme that initiates the PG synthesis including PGI2 and PGE2 which bind PTGIR and PTGER4 respectively, inducing S100A8/S100A9 (see above). Based on RNAseq data, expression of NFKB1 in THP-1 induces the expression of EBI3 (one of the top 5 HITS) (**Table 2 in S1 Table**) most likely via direct binding to the EBI3 promoter by RELB subunit. EBI3 encodes the interleukine-27 subunit beta, which can bind to IL27R and activate JAK/STAT3 pathway **(11, 12)**. In addition, NFKB1 induces the expression of PTGIR (<2 FC, RNAseq data) and by LPS **(S5 Fig)**. RELB binds directly to PTGIR promoter (***by Encode***).

Importantly, the production of CP triggered by the PTGIR or PTGER4 ligands that was observed in THP-1 cells was potentiated by the addition of LPS **(Fig 5)**. Conversely, however, stimulation of either the PTGIR or PTGER4 pathways, diminished the generation of the pro-inflammatory cytokine TNF-alpha following LPS stimulation **(Fig 6)**. While it has previously been reported that PGI2 and PGE2 have both pro- and anti-inflammatory activities in different immune cells (49), and that they can suppress the inflammatory cytokine TNF-alpha in THP-1 cells (50, 51), this is the first time that it has been observed to have the concurrent opposite effect on CP. This effect on TNF-alpha production is likely due to the inhibitory effect of STAT3 on NFkB-mediated transcription at this locus (52) (**Fig 8**). It should be noted that whereas PTGER4 is strongly expressed in monocyte/macrophages and in neutrophils from peripheral blood of healthy individuals, PTGIR expression is high in the former and lacking in the latter, suggesting that the pathway described herein is more specific to the monocyte/macrophages lineage (53).

An examination of a publicly-available eQTL dataset supports the notion that genetic control of the expression levels of the prostaglandin family members influences risk to the development of IBD. It should be noted, however, that the actual causal variants in these loci have not been elucidated. The eQTLs identified for these *PTGIR* and *PTGER4* are both associated with lower levels of expression but where *PTGER4* variant is associated with decreased risk and the *PTGIR* variant with increased risk. It is very likely that the actual direction of effect of these eQTLs is very context-dependent, for example it has previously been reported that IBD-associated variants are eQTLS in B lymphocyte-derived lymphoblastoid cell lines and are associated with higher levels of expression of *PTGER4* (17). As such, the actual levels of these receptors, their ligands and downstream effectors, as well as cellular context, may influence the resulting biological impacts.

While we did not elucidate how the other IBD genes that were found in our screen to modify CP expression in THP-1 cells, our results do suggest potential mechanisms by which they do so. First, we found that increasing the expression of *NFKB1* led to an upregulation of PTGIR expression although just shy of a two-fold threshold (FC= 1.995). *NFKB1*, which encodes a subunit of the well characterized and ubiquitous transcription factor NF-kappa-B, may also act indirectly via transcriptional control of genes such as *TNF* or *EBI3* as observed in the current screen, which would in turn stimulate the JAK2-STAT3 pathway (55, 56) leading to increased expression of *S100A8/A9* (**Fig 8 & S6 Fig**). In addition, the increased expression of *ZBTB40* induced the expression of three genes that encode key enzymes within the prostaglandin pathway (PLA2G1B, *TBXAS1* and *AKR1C2*), thus likely impacting on this pathway. Given that *ZBTB40* is a transcription factor/regulator, it likely functions by binding to some or all of the promoter regions of *S100A8, S100A9, PLA2G1B, AKR1C2* and *TBXAS1*. The mechanism by which SLC39A11 impacts on the expression of *S100A8/A9* is more speculative as it encodes a relatively uncharacterized metal ion transporter that is believed to transport zinc ions; thus its role may be linked to its control of zinc which is required to stabilize ZBTB40 and other zinc finger proteins (54). Interestingly the *SLC39A11* IBD gene led to an increase in expression of the PG E2 receptor subtype 2 (PTGER2).

Finally, together these results support an important role for the prostacyclin/prostaglandin biogenesis and signaling pathways in IBD susceptibility and pathogenesis. At the present time, there is little known about the role of prostacyclin in IBD although it has been reported that it, as well as PGE2, can regulate lymphatic and vascular functions in the intestine (57). It has also been demonstrated that PGE2 stimulation of PTGER4 in macrophages leads to the secretion of chemokine (C-X-C motif) ligand 1 (CXCL1), which in turn drives epithelial cell differentiation and proliferation from regenerating crypts, favouring mucosal healing (58). It should also be noted that CXCL1 is a chemoattractant for neutrophils and its serum level is elevated in CD patients (59, 60). Interestingly, mesenchymal stromal cells (MSCs) may be a potential source of intestinal PGE2, promoting macrophages to adopt an anti-inflammatory activation state (61). It has also been recently proposed that there is regulatory interaction between IL-10 and PGE2, that when this balance is perturbed, there is aberrant macrophage activation that contributes to IBD pathogenesis (62). Importantly, calprotectin has been shown to mediate a variety of biological functions that are key to homeostatic antimicrobial defenses but that in the chronic disease state it may exacerbate inflammation in patients with IBD (63).

## Conclusion

In conclusion, the current study provides evidence that five genes (*PTGIR, PTGER4, ZBTB40, NFKB1* and *SLC39A11*) associated with IBD susceptibility are also involved in, directly or indirectly, the control of the expression of calprotectin genes *S100A8* and *S100A9*. This work also supports that prostacyclin/prostaglandin biogenesis and signaling pathways have an important role in IBD pathogenesis.

## Methods

### Selection, cloning and lentiviral transfer of IBD candidate genes from GWAS regions

To identify the IBD gene candidates to test in our transcriptome-based screen for IBD gene functions, we focused on the 163 IBD-associated loci identified by the International IBD Genetics Consortium(12). Sixty-five genes were selected from these loci based on their endogenous expression in human cell lines and primary cells of the monocyte/macrophage lineage **(Table 1 in S1 Table)**. We cloned the open reading frames (ORFs) for these genes into a modified GATEWAY® compatible polycistronic lentiviral expression vector, pLVX-EF1a-IRES-PURO/eGFP, for expression in the THP-1 monocyte cell line model. To generate stable cell lines expressing the different candidate ORFs, THP-1 cells were transduced at least in triplicate via spinoculation. Transduction of the whole set of IBD gene candidate ORFs was performed in three batches of about 15 ORFs, as well as an empty vector control. 43 IBD-ORFs successfully cloned and transduced in THP-1 **(S1 Appendix)**.

### Preparation of the total RNA library and sequencing

Total RNA was extracted from stably transduced THP-1 cultures using the RNeasy Plus Mini kit (Qiagen) according to manufacturer’s protocol. The RNA samples were quantified, and quality controlled using an Agilent RNA 6000 Nano kit (Agilent) on an Agilent Bioanalyzer 2100 system. Samples with RNA Integrity Number (RIN) below 8 were discarded. RNA sequencing was performed at the McGill University/Genome-Québec Innovation Center. Briefly, RNA samples were transformed into barcoded DNA libraries using TruSeq Stranded mRNA library preparation kits (Illumina), which were then paired-end sequenced, generating 2×100bp reads, using an Illumina HiSeq2000 sequencer. Raw FASTQ sequences were downloaded from the platform’s server for local processing and analysis **(S1 Supporting information)**.

### Bioinformatic annotation of HITS identified in the screen

The set of genes that increased or decreased in response to the increased expression each ORF – named “HITS” - were defined as genes where the fold effect computed from the combined replicates was larger than two, and the expression was outside the expected range of variation based on the entire dataset. In order to find biological categories enriched in HITS identified in the screen, we performed enrichment analyses using the g:GOSt functional profiling tool from the online g:Profiler service (18) these analyses included gene and phenotype ontologies, biological pathways, protein database and regulatory motifs in DNA. In addition, different cis-regulatory motif analyses of the proximal promoters of these HITS have been done using PRIMA method (19) as implemented in the EXPANDER software (v8.0) **(S1 Supporting information)**.

### Validation of *PTGIR* (expression/knockdown) effect on THP-1

For the validation of the effect of *PTGIR* on THP-1, an independent set of stable cell lines expressing the *PTGIR* ORF were generated via three independent transductions with lentivirus containing the *PTGIR*-ORF or the empty lentiviral plasmid. In addition, three cell lines where the endogenous expression of *PTGIR* was knocked down via lentiviral shRNA transfer were generated. The RNA expression levels of *PTGIR, S100A8* and *S100A9* genes were quantified by qPCR. **(S1 Supporting information)**.

### Pharmacologic assessment of *PTGIR* and PTGER4 signaling pathways

THP-1 cells were centrifuged and resuspended in fresh complete media in a 24-well plate at a concentration of 10^6^ cells/mL and grown for 24 hours. The cells were then treated for 24h with different combinations of PTGIR agonist (Beraprost sodium; Sigma), PTGIR antagonist (Ro 1138452 hydrochloride; Tocris), PTGER4 agonist (CAY10684; Cayman), adenylyl cyclase activator (Forskolin; Sigma), adenylyl cyclase inhibitor (MDL12330A; Cayman), STAT3 inhibitor (Stattic; Cayman), or LPS (Sigma-Aldrich).

### Generation, characterization and testing of hiPSC-derived monocytes

Human induced pluripotent stem cells (hiPSC) have been generated by reprogramming lymphoblastoid cell lines (LCL) and then differentiated into monocytes **(S1 Supporting information)**. These monocyte cultures were immunophenotyped by flow cytometry. An enriched population of CD14+ cells was obtained via MACS purification **(S1 Fig)**. Independent hiPSC-derived monocytes obtained from different individuals were centrifuged and resuspended in fresh complete media in a 24-well plate at an average of 250,000 cells/mL and then treated for 24h with either PTGIR agonist or carrier. Levels of *S100A8/A9* were then evaluated by qPCR.

### IBD SNPs overlapping with eQTLs identified in CD14+ monocytes

We looked for overlap of SNPs implicated by IBD GWAS and eQTLs previously reported in human monocytes and macrophages. Specifically, we tested the GWAS index SNPs for the 43 genes tested in this study, as well as for *PTGER4*. As these SNPs are not necessarily causal with regard to phenotype, but very likely in linkage disequilibrium (LD) with the responsible allele, we searched for all SNPs highly correlated with these index SNPs. We therefore identified proxy SNPs (r^2^ > 0.7) in the 500 kb window for each of the IBD SNPs using the “proxy search” online tool from SNIPA-Single Nucleotide Polymorphism Annotator “(https://snipa.helmholtz-muenchen.de/snipa3/). We ran the SNPs using the GRCh37 genome assembly, the 1000 Genomes, Phase 3 v5 variant set, set the population to European and the genome annotation to Ensembl 87. We detected 1,695 proxy SNPs (**Table 6 in S1 Table**); This set of 1,695 SNPs were then examined for overlap with the cis-eQTLs (having an FDR < 0.05) identified in CD14+ human monocytes, naïve and activated by LPS at 2h and 24h or with IFN at 24h reported by Fairfax et al (20) (**Table 7 in S1 table**).

## Ethics statement

The conversion of Human lymphoblastoid cell lines (LCL) collected by the NIDDK IBD Genetics Consortium into hiPSC lines and the use of hiPSC-derived macrophage models in this study were approved by the Montreal Heart Institute Ethics Review Board.

## Acknowledgements

We would like to thank Drs. David Root (Broad Institute) and Gerald T. Nepom (Benaroya Research Institute) for their helpful discussions. We would like to thank the Sequencing team at the Génome Quebec McGill Innovation Centre for generating high quality RNA sequence data. We would like thank members of the Laboratory for Genetics and Genomic Medicine, in particular Jean Paquette, Azadeh Alikashani, Louise Thauvette, Sonia Deschenes, Geneviève Lavallée, Jessica Desjardins, and Marie-Pier Mathieux for their excellent technical support.The members of the iGenoMed Consortium at the time of this study were (in alphabetical order):

Alain Bitton^1*^, Gabrielle Boucher^2^, Guy Charron^2^, Christine Des Rosiers^2,3*^, Anik Forest^2^, Philippe Goyette^2^, Sabine Ivinson^4^, Lawrence Joseph^5*^, Rita Kohen^1^, Jean Lachaine^6*^, Sylvie Lesage^3,7*^, Megan Levings^4*^, John D. Rioux^2,3*^, Julie Thompson-Legault^2^, Luc Vachon^8^, Sophie Veilleux^9*^, Brian White-Guay^3*^. **Affiliations:** ^1^McGill University Health Centre, Montreal, Quebec; ^2^Montreal Heart Institute Research Center, Montreal, Quebec; ^3^Université de Montréal, Faculté de Médecine, Montreal; ^4^University of British Columbia, Vancouver; ^5^McGill University, Faculty of Medicine, Department of Epidemiology, Biostatistics and Occupational Health; ^6^Université de Montréal, Faculté de Pharmacie; ^7^Maisonneuve-Rosemont Hospital, Research Center, Montreal; ^8^LV Consulting, Montreal; ^9^Université de Laval, Québec. *Principal Investigators on grant # GPH-129341 with JDR as Leader and AB as co-Leader.

## Supplementary Information Captions

**S1 Table: Supplementary Tables. Table 1** Selection of genes to include in expression screen. **Table 2:** Total HITS Up/Down regulated by each ORF in THP-1 cells. **Table 3:** gProfiler enrichment analyses of the HITS identified in the screen. **Table 4:** Transcription factor binding site analysis of HITS upregulated by different IBD-ORFs by gProfiler. **Table 5** Transcription factor binding site analysis of HITS upregulated by different IBD-ORFs by PRIMA-EXPANDER. **Table 6:** Forward and reverse primer sequences 5’-3’ used in the study. **Table 7: Proxy** SNPs (r2 > 0.7) in the 500 kb window for each of the 224 IBD SNPs; **Table 8** Overlapping of the IBD proxy SNPs with the cis-eQTLs data in CD14+ monocytes.

## Supplementary Figures

**<u>S1 Fig. Generation of a hiPSC-derived monocytic model</u>. (A)** A schematic representation illustrating the different steps involved in the generation of hiPSC-derived monocyte cultures from LCLs. Briefly, LCLs are reprogrammed into hiPSC, as described by Kumar et al.(9), via nucleofection with four episomal reprogramming plasmids (pCE-hUL, pCE-hSK, pCE-hOCT3/4, and pCE-mp53DD) and selection of TRA-1-60 positive reprogrammed hiPSC colonies. LCL-derived hiPSC are then used to generate monocyte cultures using a stepwise differentiation protocol involving a conversion to colonies of myeloid progenitor cells followed by the formation of myeloblasts and then monoblast cultures. After a minimum of 16 days in culture, monoblast cultures will start releasing monocytes in the culture supernatant; typically, an average of 500,000 monocytes can be harvested every 3-4 days for use in specific experiments following characterization by flow cytometry. Conversion of LCLs to hiPSC is validated by **(B)** the increased expression of pluripotency markers such as POU5F1 and NANOG (by qPCR), as well as **(C)** the expression of TRA-1-60 by hiPSC colonies (by immunofluorescence). **(D)** The pluripotent potential of the hiPSC lines is then evaluated through their ability to convert into the three germ layers (by immunofluorescence). (E) The harvested monocytes are then evaluated first for the loss of the pluripotence marker TRA-1-60 (dashed histogram: unstained cells, grey histogram: isotype control, blue histogram: anti-TRA-1-60 staining) and morphologically by H&E staining. **(F)** CD14+ cells (typically, >90% of harvested cells) are then further validated via FACS analysis of the CD11b and CD45 cell-surface markers. Gating was determined using isotype control staining.

**<u>S2 Fig. Impact of ZBTB40 expression on S100A8/9 genes expression in THP1</u>**. Relative mRNA expression levels of (**A**) endogenous ZBTB40 and of **(B)** ORF-driven (optimized) ZBTB40, as well as **(C**) fold change in mRNA expression of S100A8 and S100A9 were evaluated in THP-1 cell lines transduced with lentiviruses containing either empty plasmid or the ZBTB40 ORF. Each bar is the mean of 3 samples from 3 different infections ±SEM. RNA expression was determined using qPCR in duplicate and was normalized to β-actin mRNA level. \*\*\**P* < .001. For primer sequences used to quantify endogenous ZBTB40 and ZBTB40 ORF (optimized) expression levels **see table 6 in S1 Table**.

**<u>S3 Fig. Impact of PTGIR expression on the CP (S100A8/9 dimer) protein levels in THP1</u>**. The total intracellular levels **(A)** and extracellular concentration **(B)** of CP protein was determined by ELISA from 5×10^5^ cells from each cell line (transduced with either empty plasmid or by PTGIR ORF) cultured in 0.5ml complete medium for 24 h. Cell pellets and culture supernatants were collected for the evaluation of total intracellular CP levels and extracellular CP concentration respectively. Each bar is the mean of 3 samples from 3 different infections ±SEM. \*\**P* < .01. **(C)** CP secreted concentration measured by ELISA 24 h after THP-1 incubation with 10^−5^ M PTGIR agonist (Beraprost Sodium). Each bar is the mean of 4 samples from 4 different experiment ±SEM. \**P* < .05

**<u>S4 Fig. Identification of TFBS in the promoter regions of S100A8 and S100A9</u>**.

Promoter regions (1000 bps upstream of the transcription start site) of S100A8 and S100A9 were evaluated for DNase clusters and transcription factor ChIP-seq Clusters (340 factors, 129 cell types) from ENCODE 3 (data version: ENCODE 3 Nov 2018); UCSC Genome Browser on Human Feb. 2009 (GRCh37/hg19) Assembly. Red arrow indicates the binding site of STAT3.

**<u>S5 Fig. Impact of LPS on PTGIR expression in THP-1</u>**. Relative mRNA expression level of PTGIR was evaluated after incubating THP-1 for 24 hours with or without 0.2 ug/ml of LPS. Each bar is the mean of 3 samples from 3 different experiments ±SEM. RNA expression was determined using qPCR in duplicate and was normalized to β-actin mRNA level. \*\**P*<.01 (Student’s *t*-test paired).

**<u>S6 Fig. Impact of TNF-alpha cytokine on S100A8/9 genes expression in THP-1</u>**.

Relative mRNA expression levels of CP genes were evaluated after incubating THP-1 with either DMSO or 20ng/ml of TNFa cytokine. RNA expression was determined using qPCR in duplicate and was normalized to β-actin mRNA level. Each bar is the mean of 2 samples from 2 experiment ±SEM.

**<u>S7 Fig. The impact of different concentrations of PTGIR agonist on CP genes</u>**. Relative mRNA expression levels of S100A8 and S100A9 genes were evaluated after incubating parental THP-1 cells with either DMSO or increasing doses of PTGIR agonist (Beraprost Sodium) for 24h. RNA expression was determined using qPCR in duplicate and was normalized to β-actin mRNA level. Each bar is the average of PCR duplicates values of 1 sample from **1** experiment ±SEM.

**<u>S8 Fig. The impact of PTGIR agonist on CP genes at different time</u>**. Relative mRNA expression levels of S100A8 and S100A9 genes were evaluated after incubating parental THP-1 cells with either DMSO or 1×10^−5^ M of PTGIR agonist (Beraprost) for 24h and 48h. RNA expression was determined using qPCR in duplicate and was normalized to HPRT mRNA level. Each bar is the average of PCR duplicates values of 1 sample from 1 experiment ±SEM.

**<u>S9 Fig. The impact of different concentrations of PTGIR antagonist on CP genes expression in THP1</u>**. Relative mRNA expression levels of S100A8 and S100A9 genes were evaluated after incubating a THP-1 cell line stably transduced and expressing the PTGIR ORF with either DMSO or increasing doses of the PTGIR antagonist (Ro 1138452) for 24 h. RNA expression was determined using qPCR in duplicate and was normalized to β-actin mRNA level. Each bar is the average of PCR duplicates values of 1 sample from 1 experiment ±SEM.

**<u>S1 Appendix</u>: Impact of the expression of all 43 IBD gene candidate ORFs on the transcriptome of THP-1 cells**. Graphs illustrating the impact observed on the transcriptome of THP-1 cells following the expression of each ORF (43 ORFS). In the first four (ZBTB40, SLC39A11, NFKB1, PTGIR), the S100A8 and S100A9 genes are labeled. For the description of graphs see *Methods section*.

**<u>S1 Supporting information</u>: Supplementary Methods**.

